# BackWards – Unveiling the Brain’s Topographic Organization of Paraspinal Sensory Input

**DOI:** 10.1101/2022.08.05.502912

**Authors:** Alexandros Guekos, David M Cole, Monika Dörig, Philipp Stämpfli, Louis Schibli, Philipp Schuetz, Petra Schweinhardt, Michael L Meier

## Abstract

Cortical reorganization and its potential pathological significance are being increasingly studied in musculoskeletal disorders such as chronic low back pain (CLBP) patients. However, detailed sensory-topographic maps of the human back are lacking, and a baseline characterization of such representations, reflecting the somatosensory organization of the healthy back, is needed before exploring potential sensory map reorganization. To this end, a novel pneumatic vibrotactile stimulation method was used to stimulate paraspinal sensory afferents, while studying their cortical representations in unprecedented detail. In 41 young healthy participants, vibrotactile stimulations at 20 Hz and 80 Hz were applied bilaterally at nine locations along the thoracolumbar axis while functional magnetic resonance imaging (fMRI) was performed. Model-based whole-brain searchlight representational similarity analysis (RSA) was used to investigate the organizational structure of brain activity patterns evoked by thoracolumbar sensory inputs. A model based on segmental distances best explained the similarity structure of brain activity patterns that were located in different areas of sensorimotor cortices, including the primary somatosensory and motor cortices and parts of the superior parietal cortex, suggesting that these brain areas process sensory input from the back in a “dermatomal” manner. The current findings provide a sound basis for testing the “cortical map reorganization theory” and its pathological relevance in CLBP.

**Highlights:**

- Fine-grained cortical activation patterns of paraspinal vibrotactile sensory input were obtained using whole-brain representational similarity analysis.
- The patterns were well explained using a model reflecting segmental distances along the thoracolumbar axis.
- The current results provide a solid basis for revisiting the “cortical map reorganization theory” and its pathological significance in chronic low back pain.

## 1. Introduction

In the vast majority of patients with chronic low back pain (CLBP), no clear-cut physiological or anatomical pathology can be identified (Maher et al., 2017, Maher and Ferreira, 2022). Mounting evidence suggests that CLBP is linked to cortical alterations (Brumagne et al., 2019, Van Dieën et al., 2017, Vittersø et al., 2022). Reorganization of the primary somatosensory cortex (S1) map, i.e., altered somatotopic representation of the back, is considered a prominent example of maladaptive cortical alterations in CLBP (Brumagne et al., 2019, Van Dieën et al., 2017, Wand et al., 2011, Jenkins et al., 2022). Yet, it is difficult to position findings of S1 map reorganization in CLBP within a coherent narrative, because only very little is known about the somatosensory representation of the human back in general. Detailed somatosensory maps of the back are lacking. Specifically, there is no knowledge of the somatosensory representational organization of paraspinal sensory input along the thoracolumbar axis. A baseline characterization of such representations, reflecting the somatosensory organization of the healthy back, is needed as a basis for further investigations of potential sensory map reorganization.

A suitable approach for cortical mapping of sensory representations from different body parts is based on vibrotactile stimulation. Paradigms using vibrotactile stimuli are an established method for assessing the cortical topographic representations of different body parts (Goossens et al., 2016, Harrington and Downs III, 2001, Kim et al., 2016, Schellekens et al., 2021, Willoughby et al., 2021). Mechanical afferent inputs, including vibration, are transformed via signal transduction into electrical signals and relayed via primary afferents to the dorsal column-medial lemniscus pathway before being transmitted via the brainstem through the thalamus to the sensorimotor cortices (Kandel et al., 2013). Vibrotactile stimulation at specific frequencies can be used to assess cortical responses to the activation of different mechanoreceptors (Harrington and Downs III, 2001, Kim et al., 2016). More specifically, it has been shown that certain mechanoreceptors, such as Meissner corpuscles or Merkel disks, are activated at lower frequencies and deeper located Pacinian corpuscles or muscle spindles at higher frequencies (Avanzino et al., 2014, Schellekens et al., 2021, Weerakkody et al., 2007).

To apply vibrotactile stimulation to the back, a novel apparatus, the pneumatic vibration device (pneuVID), was recently developed (Cole et al., 2022, Schibli et al., 2021). It can apply controlled vibrotactile stimuli across different segments of the thoracolumbar axis at specific frequencies, and has been validated for use in magnetic resonance imaging (MRI) environments for functional (fMRI) data acquisition (Schibli et al., 2021). Furthermore, the usefulness of pneuVID in assessing neural representations of the human back has recently been tested using univariate statistical neuroimaging techniques (Cole et al., 2022). Univariate fMRI analysis based on net blood oxygenation level-dependent activity levels is a first step towards exploring cortical representations of different body parts. However, it cannot easily provide a deeper understanding of cortical (topographic) organization (Makin and Flor, 2020, Popov et al., 2018); for example, in the current context, revealing the organizational structure of cortical representations of paraspinal sensory inputs along the thoracolumbar axis.

Multivoxel pattern analysis techniques such as representational similarity analysis (RSA) can reveal how neural information is architectonically represented and allows for assessing and relating measured neural activity to computational and behavioral models (Kriegeskorte et al., 2008, Popal et al., 2019, Ejaz et al., 2015, Dimsdale Zucker and Ranganath, 2018, Dimsdale-Zucker et al., 2018). Pattern information is stored in a representational dissimilarity matrix (RDM), the cells of which quantify the spatial dissimilarity in neural activation for each pair of experimental conditions, thereby reflecting the (spatial) organizational structure of brain activity patterns. This so-called brain RDM can then be tested for the strength of agreement with the RDM of an explanatory model (model RDM).

Using RSA, this study aimed to provide novel insights into the cortical sensory-topographic representations of the back, specifically of paraspinal vibrotactile stimulation across different thoracolumbar segments at two specific frequencies (20 and 80Hz). A whole-brain searchlight analysis (Nili et al., 2014) was performed to test for brain RDMs with significant agreement with model RDMs that reflect different degrees of brain pattern distinguishability w.r.t. thoracolumbar segmental sensory input. The best-fitting model was hypothesized to yield informative (model-consistent) brain activity patterns primarily located in somatosensory cortices, thereby revealing brain areas that share specific neural information related to segmental paraspinal sensory input.

## 2. Material and Methods

### 2.1. General Study Design

The study followed a within-subject experimental design including one MRI session with vibrotactile stimulation of the back. Study approval was obtained from the ethics board of the Canton of Zurich. The study adhered to the principles of the Declaration of Helsinki.

### 2.2. Participants

All participants (N=41, 25 women) were healthy pain-free adults between 18 and 39 years of age (29.6 *±* 5.2 years) and a body mass index below 30 kg/m^2^ (22.4 *±* 2.6). Exclusion criteria were the presence of any back pain within the past three months, any history of chronic pain, prior spine, foot, or ankle surgery, any history of psychiatric or neurological disorders, excessive consumption of alcohol (*≥*2 standard glasses/d for women, *≥*4 standard glasses/d for men), other intoxicants, or analgesics within the past 24 hours, impairments of the motor system, and MRI contraindications.

### 2.3. Testing Procedure

#### 2.3.1. Participant Setup

Nine spinous processes, *T*3, *T*5, *T*7, *T*9, *T*11, *L*1, *L*3, *L*5, and *S*1 (*T*=thoracic, *L*=lumbar, *S*=sacral), were identified on the back of every participant by an experienced physiotherapist for subsequent attachment of the pneuVID units. For the duration of the experiment, participants were lying in a supine position on the bed of the MRI scanner with their eyes open.

#### 2.3.2. Pneumatic Vibration Device (pneuVID)

Full details on the pneuVID methodology and its integration into the MRI environment have previously been outlined (Schibli et al., 2021, Cole et al., 2022). In short, pneuVID units were attached bilaterally to the identified spinous processes over the erector spinae muscles. A valve box delivered pulses of compressed air to the pneuVID units, and a control module (Raspberry Pi, Cambridge, UK) connected the stimulation setup to the MR system. Prior to the fMRI data acquisition, a sensory testing session was performed to ensure that participants were able to distinguish between the sensations evoked by the two stimulation frequencies and to rule out lateralization of sensory perception along the thoracolumbar axis.

#### 2.3.3. Stimulation Protocol

Stimulation was applied at frequencies of 20 and 80Hz. Amplitudes ranged from 0.5 to 1mm because a fixed amplitude could not be guaranteed due to material properties of the vibrational units (Schibli et al., 2021). Each stimulus had a duration of five seconds, with a jittered interstimulus interval between 4 and 7s. Two experimental runs were performed with six stimuli per location and frequency, that is, 108 stimuli per run (9 locations *·* 2 frequencies *·* 6 stimuli = 108 stimuli). The stimulation order was pseudorandomized between participants, with the restriction of a maximum of three consecutive stimuli at the same location or frequency. The air pressure of the delivered pulses was maintained constant at 1.5bar during the experiment.

The two stimulation frequencies of 20 and 80Hz were chosen because they have been suggested to activate different tactile and proprioceptive afferents of superficial and deep mechanoreceptors (Avanzino et al., 2014, Roll et al., 1989, Schellekens et al., 2021). Meissner corpuscles are targeted at approximately 20Hz, whereas frequencies around 80Hz have been shown to be a potent stimulus for activating muscle spindles, which are the main proprioceptors in the human body (Biggio et al., 2021, Proske and Gandevia, 2012). However, it should be noted that these frequency-mechanoreceptor relationships are not exclusive, as for example Pacinian corpuscles may also be co-stimulated during 80Hz stimulation (bandwidth between 50 and 400Hz) and have been shown to transmit proprioceptive information through skin stretch (Chung et al., 2013, LaMotte and Mountcastle, 1975, Talbot et al., 1968, Weerakkody et al., 2007). Hence, a perfectly isolated examination of the stimulated mechanoreceptor type is not realistic in the current experimental context.

#### 2.3.4. Neuroimaging Data Acquisition

Data acquisition was performed on a Philips Achieva 3T MRI scanner with a head coil of 32 channels (Philips, Best, The Netherlands). Each stimulation run lasted 22.5 minutes. Per run, 750 volumes of echo-planar imaging (EPI) were acquired with T2*-weighting (TR=1.8s; TE=34ms; flip angle=70*^◦^*; 54 interleaved ascending axial slices with multiband factor 3; in-plane resolution=1.72*·*1.72mm^2^; slice thickness=2.0mm with no slice gap; sensitivity encoding factor=1.4; EPI factor=87). Between the two runs, a T1-weighted anatomical scan was acquired (sequence=MPRAGE; TR=6.6ms; TE=3.1ms; flip angle=9*^◦^*; field of view=230*·*226*·*274; voxel size=1.0*·*1.0*·*1.2mm^3^; turbo field echo factor=203) resulting in a total scan duration of approximately 50 minutes per participant.

### 2.4. Representational Similarity Analysis (RSA)

The Python Representational Similarity Analysis toolbox (rsatoolbox v.0.0.4) (Kriegeskorte et al., 2021) was used to implement RSA. It was carried out on Jupyter Notebooks v1.0.0 (Jupyter, 2021) with Python v3.8.10 (Python Software Foundation, Delaware, USA) and NiBabel v3.2.2 (Brett et al., 2022), Nilearn v0.9.0 (Abraham et al., 2014), NLTools v0.4.5 (Chang et al., 2021), Pandas v1.4.1 (McKinney and the Pandas Development Team, 2022), and Seaborn v0.11.2 (Waskom, 2021). RSA was run in a supercomputing environment (ScienceCloud, University of Zurich, Switzerland).

#### 2.4.1. Data Pre-Processing

Pre-processing of the neuroimaging data followed a previously published pipeline (Cole et al., 2022).

For every participant, mean activation contrasts (Z-transformed t-value maps) normalized to the Montreal Neurological Institute (MNI) standard space (2mm isotropic voxels) were obtained for each of the nine conditions, corresponding to the nine paraspinal locations (see supplementary figure [S2] in (Cole et al., 2022) for individual brain activation maps). Using FSL FEAT, first-level analysis outputs were assessed on the participant level to test for mean representation of contrast parameter estimates per run. Separate contrasts were defined for the two stimulation frequencies, i.e. for 20Hz and 80Hz compared to baseline (unsmoothed data).

#### 2.4.2. Searchlight Approach

In a RSA searchlight analysis, a small volume is moved over a pre-defined brain region (Dimsdale-Zucker and Ranganath, 2018). In the present study, a whole-brain spherical searchlight analysis was performed with a searchlight diameter of 5 voxels (Kriegeskorte et al., 2006, Etzel et al., 2013), resulting in a total of 155,815 searchlights.

#### 2.4.3. Representational Dissimilarity Matrices of Neural Activation (Brain RDMs)

For every participant, a 9*·*9 brain RDM was created for each contrast. A single RDM contains the dissimilarity information of neural activation patterns for every pair of experimental conditions, i.e., for every pair of stimulated paraspinal locations. By way of construction, every brain RDM is symmetrical with a diagonal of zeros regardless of the dissimilarity measure chosen. In this study, the dissimilarity was quantified using the crossvalidated Mahalanobis distance (Crossnobis), which gives an estimate of the geometric distance between two activation patterns taking into account noise covariance and using data partitioning for crossvalidation (Kriegeskorte et al., 2006, Walther et al., 2016). Crossnobis has been shown to be the most reliable dissimilarity measure for multivoxel pattern analysis (Walther et al., 2016).

#### 2.4.4. Model Representational Dissimilarity Matrices (Model RDMs)

Figure 1 shows the schematics and the entries of the three experimental model RDMs that were used for RSA. The three models represent different degrees of spatial distinguishability of brain activity patterns evoked by paraspinal sensory inputs across the thoracolumbar segments.

First, the *segmental model* reflects the organization of brain activity patterns that resemble the organizational principles of human dermatome maps of the back (Downs and Laporte, 2011). Paraspinal sensory input from neighbouring segments should result in more similar patterns than that from segments located farther apart; that is, increasing anatomical distance should lead to higher dissimilarity in patterns. The dissimilarity entries in the segmental model RDM thus increase by 0.1 with distance (except for *L*5 to *S*1, which was a priori set to 0.05, owing to the comparatively shorter physical distance between these segments on the back).

Second, the *simple model* only distinguishes between the thoracic (*T*3-*T*11) and lumbosacral (*L*1-*S*1) regions; therefore, it categorially differentiates between upper and lower back paraspinal sensory input. As such, the simple model RDM contains zeros (i.e., total similarity) for pairs within the same region and ones (i.e., total dissimilarity) for pairs within distinct regions.

Third, the *random model* corresponds to an arbitrary distribution of brain activity patterns induced by segmental sensory input. This model contained no explanatory information and was used as a control in the model comparison.

**Figure 1:**
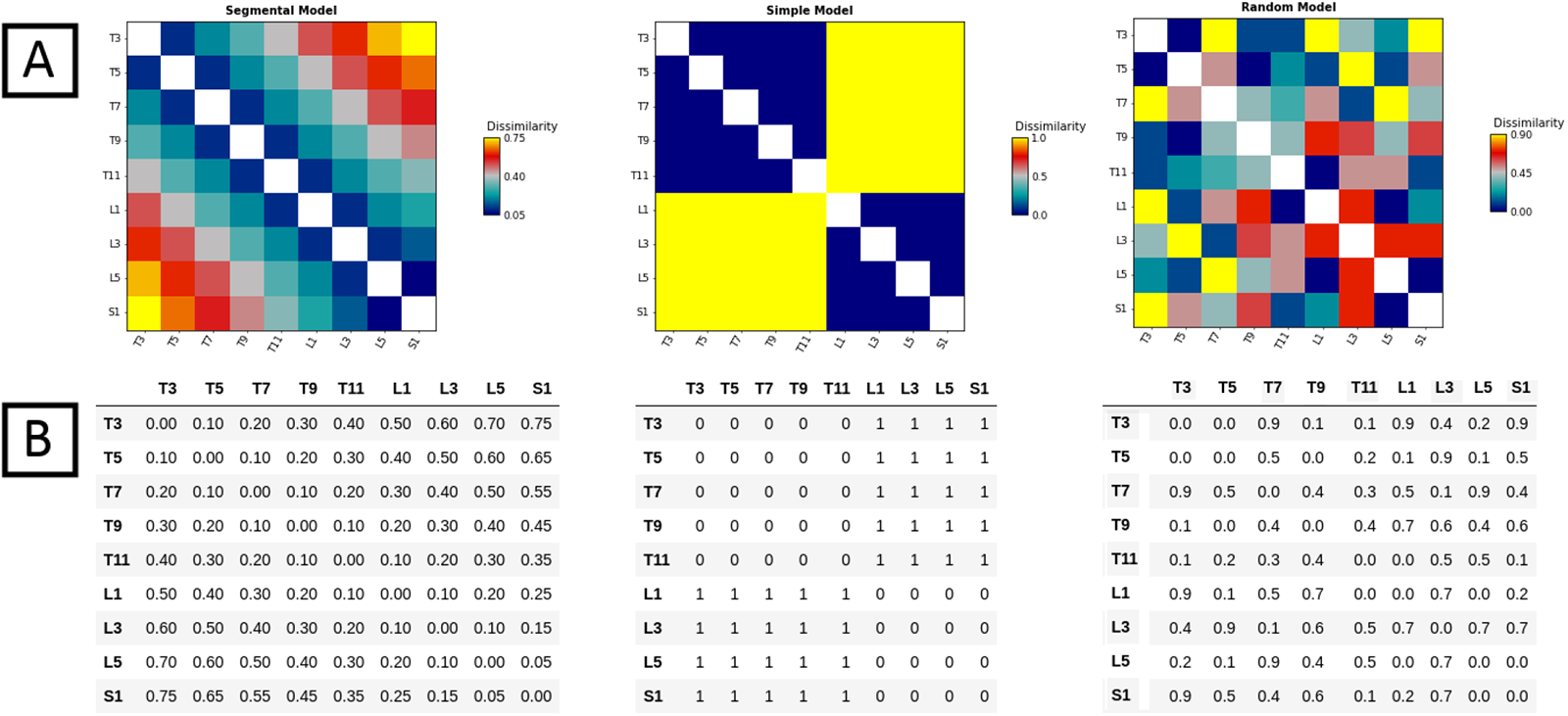
Representational Dissimilarity Matrices for the three experimental models (model RDMs). A: Colour-coded schematics of the model RDMs (*segmental model*, *simple model*, *random model*) with vertical bars indicating the dissimilarity colour scale (0=total similarity; 1=total dissimilarity). B: Numerical entries of the three model RDMs. Lines and columns are labelled according to the nine stimulated spinous processes *T*3-*S*1 (*T*=thoracic, *L*=lumbar, *S*=sacral).

### 2.5. Representational Similarity Analysis

To compare brain RDMs to model RDMs, Kendall’s rank correlation coefficient (Kendall’s *τ_a_*) was used (Kendall, 1970), a robust measure for models predicting tied ranks (Nili et al., 2014, Popal et al., 2019). For every subject, this resulted in a vector of correlation coefficients whose length corresponds to the number of searchlights. The coefficients were Fisher-z transformed (Walker, 2003).

Using FSL Randomise (FSL v5.0.9, FMRIB Software Library, University of Oxford, UK), to compute the p-value, 10,000 permutations were performed to obtain a null distribution of z scores. Threshold-free cluster enhancement (TFCE) was then applied to group-level cluster identification, allowing for a family-wise error (FWE) below 5% (Smith and Nichols, 2009). Between-model comparisons were performed based on difference maps created by subtracting one of the two respective z-maps from the other for every subject, before applying a one-sample t-test and TFCE with FWE correction.

The analyses were performed separately for 20 and 80Hz stimulations. Additionally, the models were tested on difference maps contrasting the 20 and 80Hz conditions.

## 3. Results

### 3.1. Segmental Model

The *segmental model* demonstrated significant agreement with the organizational structure of brain activity patterns located in medial parts of the S1, consistent with the assumed location of the back between the representations of the hip and shoulder according to the sensory homunculus (Penfield, 1947), superior parietal cortex (SPL), and M1 regions for both, 20 and 80Hz conditions (pFWE < 0.05, figure 2, table 1). At 20Hz, significant model-consistent brain activity patterns were restricted to two small, right-lateralized clusters, mostly within the S1 and the SPL. These clusters were primarily located within Brodmann area (BA) BA5 and, to a lesser extent, BA7, BA2, BA1, BA3b, BA4p, and BA6. At 80Hz, significant model-consistent brain activity patterns were observed in bilateral sets of brain areas within the SPL, S1, and M1. Again, the clusters were primarily located within BA5 and extended into BA7, BA2, BA1, BA3a, BA3b, BA4a, BA4p, and BA6.

**Figure 2:**
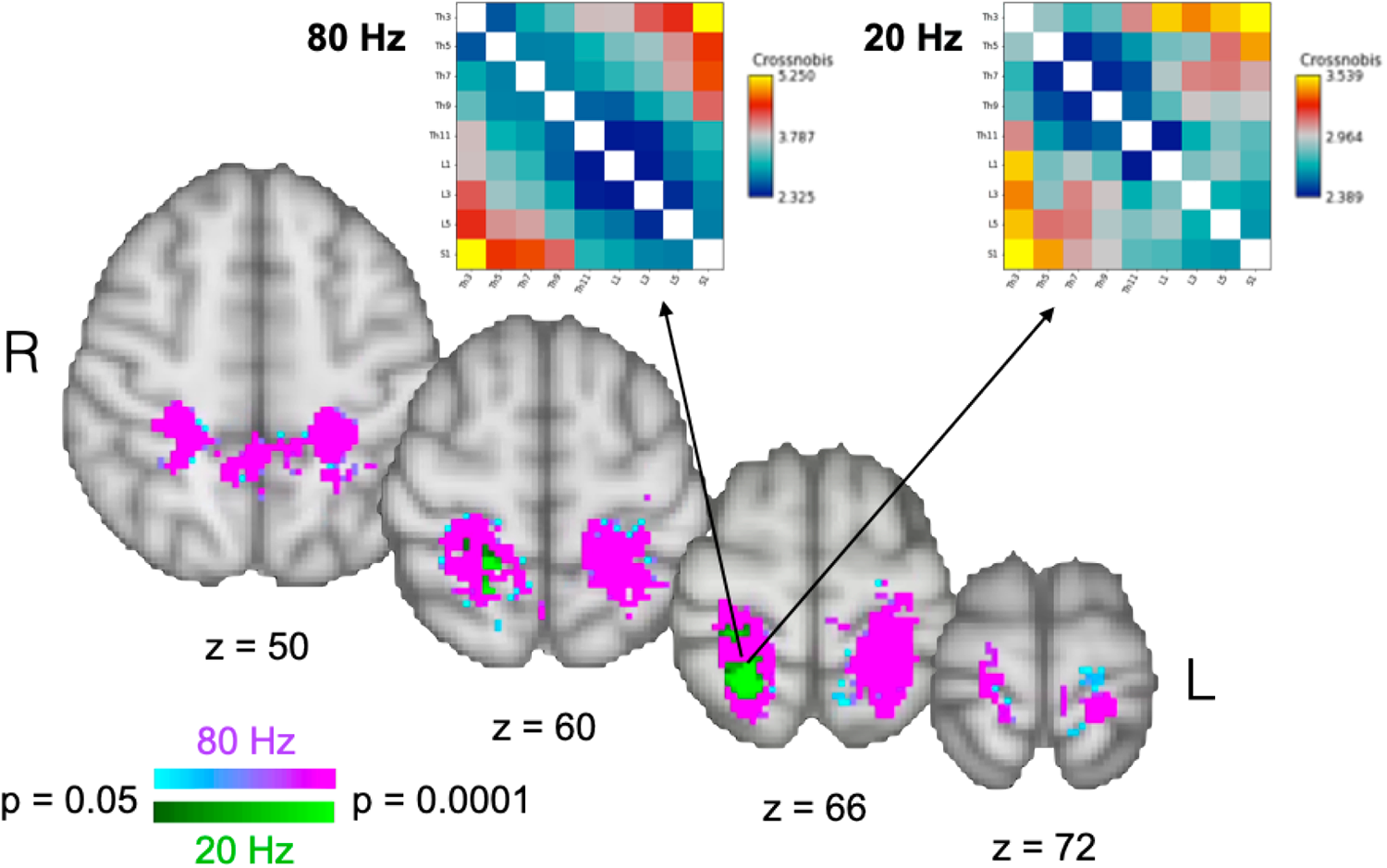
Brain areas showing significant agreement (representational similarity) of brain activity patterns to the *segmental model* representational dissimilarity matrix (RDM). Horizontal colour bars represent significant p-values following permutation testing for the 80Hz (cyan-magenta) and 20Hz (dark green-light green) conditions. Only clusters that survived threshold-free cluster enhancement with family-wise error correction (P < 0.05) are shown. Statistical maps were overlaid on representative axial slices in the z-plane of the background image of the T1 template in the Montreal Neurological Institute (MNI152) standard space. R=right hemisphere. L=left hemisphere. Arrows indicate the best brain RDM fits to the *segmental model* RDM for the 20 and 80Hz conditions (see Table 1 for the exact searchlight centers (peak coordinates)). Lines and columns of the brain RDMs are labelled according to the nine stimulated spinous processes *T*3-*S*1 (*T* =thoracic, *L*=lumbar, *S*=sacral). The vertical colour scales represent the degree of dissimilarity, as quantified by the crossnobis distance. L=left, R=right hemisphere.

**Table 1:** Cluster information for identified brain regions using the *segmental model*.

| Frequency | Number of voxels | Peak p-value | Peak x | Peak y | Peak z | COG x | COG y | COG z | Brain regions |
| --- | --- | --- | --- | --- | --- | --- | --- | --- | --- |
| 20Hz | 155 | <0.0001 | 20 | -52 | 64 | 22 | -44 | 64 | R and L: BA5, extending into BA7, BA2, BA1, BA3b, BA4p, BA6 |
| 80Hz | 3039 | <0.0001 | 28 | -36 | 46 | 0 | -40 | 60 | R and L: BA5, extending into BA7, BA2, BA1, BA3a, BA3b, BA4a, BA4p, BA6 |

Clusters of voxels identified using the *segmental model* representational dissimilarity matrix (based on a 5-voxels diameter searchlight, 10,000 permutations with threshold-free cluster enhancement and family-wise error ( $p < 0.05$ ) correction. For every cluster, the locations of the peak p-value (searchlight centres) and the centre of gravity (COG) are given. R=right hemisphere. L=left hemisphere. BA=Brodman area.

### 3.2. Simple Model

The *simple model* was significantly explanatory for the 80Hz condition only (pFWE < 0.05, figure 3, table 2). Significant model-consistent brain activity patterns were observed in distinct lateralized regions within the S1, M1, and SPL. In the right hemisphere, at the midline, and in the left hemisphere, these clusters were primarily located within BA5, BA7, and BA2.

**Table 2:** Cluster information for identified brain regions using the *simple model*.

| Frequency | Number of voxels | Peak p-value | Peak x | Peak y | Peak z | COG x | COG y | COG z | Brain regions |
| --- | --- | --- | --- | --- | --- | --- | --- | --- | --- |
| 80Hz | 932 | <0.0001 | 22 | -40 | 50 | 20 | -42 | 60 | R and M: BA5, BA7, BA2 |
|  | 578 | <0.0001 | -18 | -36 | 52 | -22 | -42 | 62 | L: BA5, BA7, BA2 |

Clusters of voxels identified using the *simple model* representational dissimilarity matrix (based on a 5-voxels diameter searchlight, 10,000 permutations with threshold-free cluster enhancement and family-wise error ( $p < 0.05$ ) correction. For every cluster, the locations of the peak p-value and the centre of gravity (COG) are given. R=Right hemisphere, L=left hemisphere, M=midline. BA=Brodman area.

### 3.3. Random Model

The random model searchlight analysis did not yield any brain activity patterns consistent with the model for either frequency condition.

**Figure 3:**
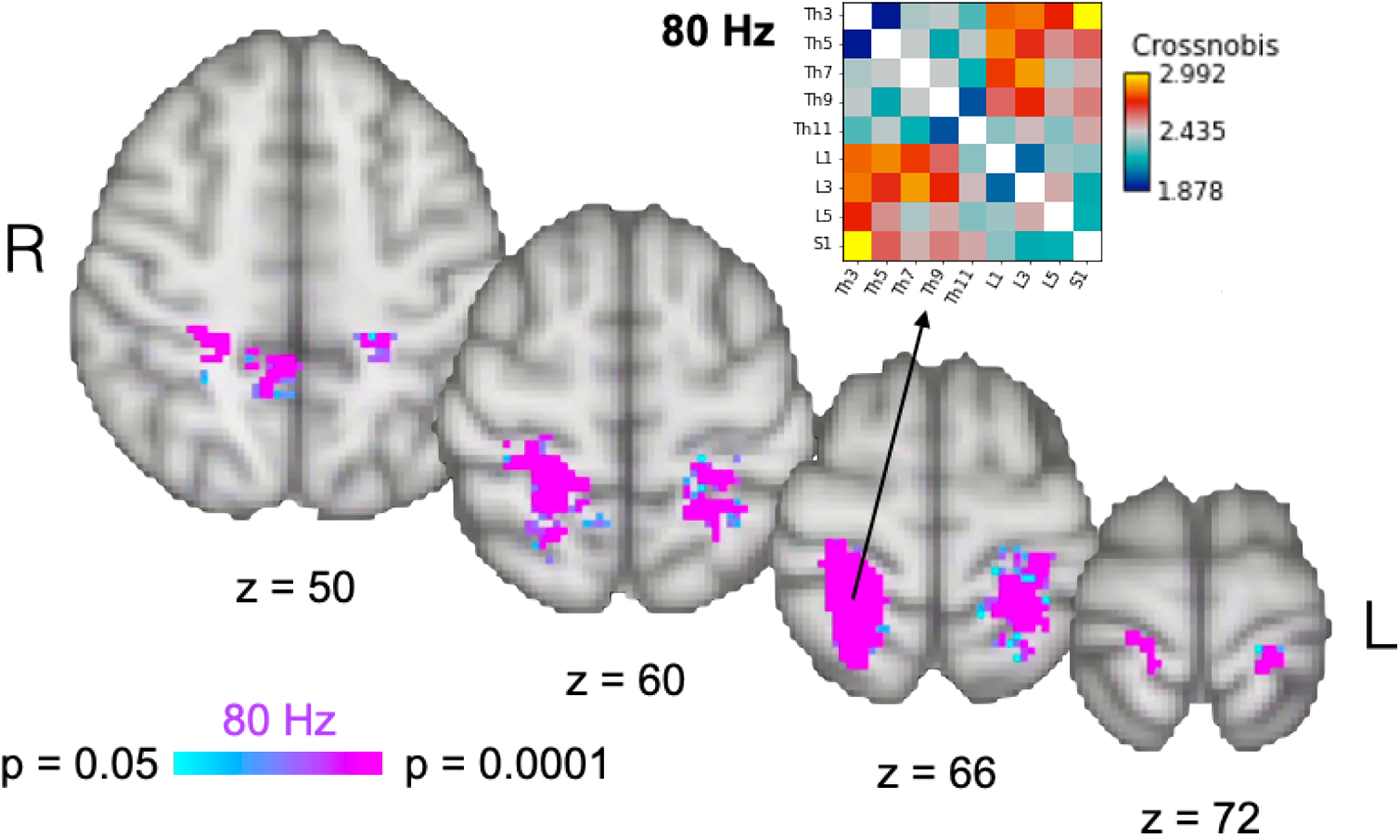
Brain areas demonstrating significant representational similarity of brain activity patterns to the *simple model* representational dissimilarity matrix (RDM). Horizontal colour bars represent significant p-values following permutation testing for 80Hz (cyan-magenta) vibrotactile stimulation conditions. Only clusters surviving threshold-free cluster enhancement with a correction for family-wise error of p<0.05 are reported. Statistical maps are overlaid on representative axial slices in the z-plane of a background image of a T1 template in Montreal Neurological Institute (MNI152) standard space. R=right hemisphere. L=left hemisphere. The arrow indicates the best group-level brain RDM fit to the *simple model* at 80Hz. Lines and columns of the brain RDM are labelled according to the nine stimulated spinous processes *T*3-*S*1 (T=thoracic, L=lumbar, S=sacral). The vertical colour scales represent the degree of dissimilarity, as quantified by the crossnobis distance. L=left, R=right hemisphere.

### 3.4. Between-model Comparisons

Comparison between models yielded a superiority of the *segmental model* in explaining the organizational structure of sensorimotor brain activity patterns evoked by 80Hz, but not by 20Hz, stimulation compared to the other two models (pFWE < 0.05, figure 4A-B and table 3). Furthermore, the *segmental model* agreement was significantly stronger for the 80 compared to 20Hz condition (pFWE < 0.05, figure 4C and table 3).

**Figure 4:**
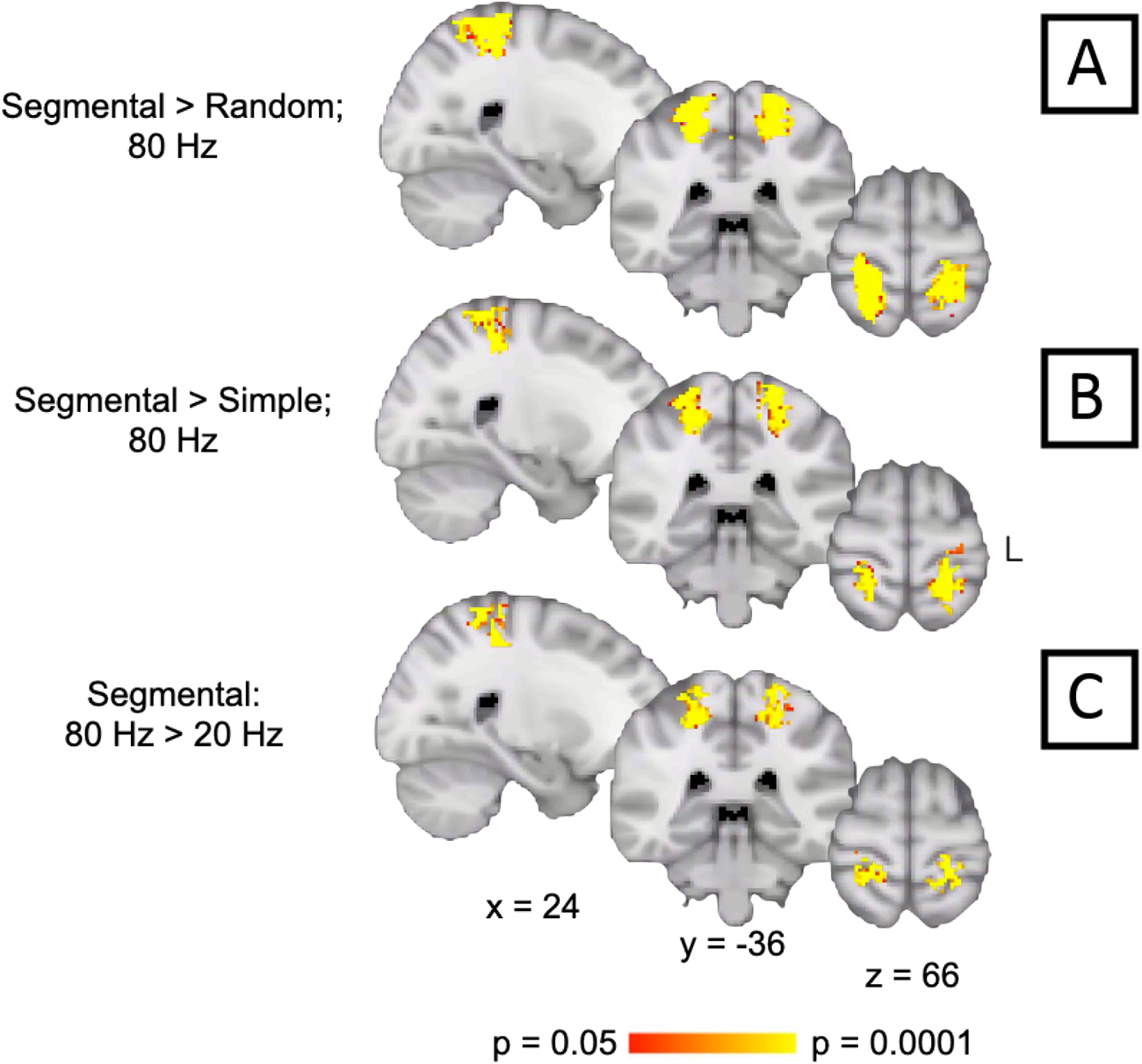
A and B: Brain activity patterns that demonstrated significantly better agreement with the *segmental model* compared to the *random model* and the *simple model* (80Hz condition). C: Brain activity patterns during 80Hz stimulation demonstrated significantly better agreement with the *segmental model* compared to 20Hz stimulation. Horizontal colour bars represent significant p-values following permutation testing (N=10,000). Only clusters surviving threshold-free cluster enhancement with a correction for family-wise error of p<0.05 are shown. Statistical maps are overlaid on three representative slices in the sagittal (x), coronal (y), and axial (z) planes of a background image of a T1 template in Montreal Neurological Institute (MNI152) standard space. L=left hemisphere.

**Table 3:** Cluster information for between-model comparisons.

| Model comparison | Number of voxels | Peak p value | Max x | Max y | Max z | COG x | COG y | COG z | COG location |
| --- | --- | --- | --- | --- | --- | --- | --- | --- | --- |
| Segmental > random; 80Hz | 1082 | <0.0001 | 24 | -36 | 46 | 20 | -42 | 60 | BA2 R |
|  | 867 | <0.0001 | -20 | -34 | 48 | -22 | -42 | 60 | BA2 L |
| Segmental > simple; 80Hz | 737 | <0.0001 | -34 | -46 | 50 | -22 | -40 | 60 | BA2 L |
|  | 407 | <0.0001 | 26 | -36 | 46 | 24 | -38 | 60 | BA2 R |
| Segmental: 80Hz > 20Hz | 453 | <0.0001 | 28 | -30 | 48 | 20 | -40 | 58 | BA2 R |
|  | 398 | <0.0001 | -20 | -36 | 48 | -22 | -38 | 60 | BA2 L |

Clusters of voxels for brain activity patterns that demonstrated significantly better agreement between models. Max = maximum; COG = centre of gravity; segmental = *segmental model*; random = *random model*; simple = *simple model*; BA = Brodmann area; R = right; L = left.

## 4. Discussion

The present study aimed at establishing sensory-topographic maps of the human back by identifying brain areas that carry distinguishable pattern information w.r.t. the thoracolumbar segmental processing of paraspinal sensory inputs. To this end, a previously validated stimulation apparatus and protocol (pneuVID) (Cole et al., 2022, Schibli et al., 2021) were used in combination with RSA. As hypothesized, the whole-brain and model-driven searchlight RSA revealed informative brain activity patterns mainly in the somatosensory cortices, with some striking differences in terms of applied model RDMs and frequency conditions. Different frequencies were chosen to explore the contributions to brain activity patterns potentially resulting from different mechanoreceptor stimulations. In general, the *segmental model* best explained the brain activity patterns in distinct areas of somatosensory cortices, indicating that these brain areas process sensory inputs from the back in a “dermatomal-like” fashion and that the applied fMRI protocol is sufficiently sensitive to reveal the corresponding pattern information. This is in line with the general observation that the spatial relationship between receptors on the body’s surface is maintained along the body-brain axis, reflected by topographically organized representations in the sensory homunculus (Puckett et al., 2020, Schott, 1993, Yamada et al., 2007). In addition, the current results indicate that this back-specific topographic pattern is shared across multiple brain regions, including the pre- and post-central gyri as well as parts of the SPL, a higher level integrative brain region that has also been shown to process topographically organized information (Liu et al., 2021). This is the first evidence of fine-grained cortical topographic information of the back, which has previously been demonstrated for other body parts such as the fingers (Besle et al., 2014, Kolasinski et al., 2016, Martuzzi et al., 2014).

### 4.1. Differences between Low- and High-Frequency Stimulation

The cortical mapping obtained from correlating brain and model RDMs showed striking differences between informative activation patterns resulting from low (20Hz) compared to high frequency (80Hz) vibrotactile stimulation on the back. The organizational structure of brain activity patterns evoked by 80Hz stimulation was well captured by the *segmental model*, which also was the best model for explaining the structure of activation patterns during 20Hz stimulation. The simple model showed significant agreement with the structure of brain activity patterns evoked by 80Hz stimulation only. Differences in the physiological characteristics of the activated mechanoreceptors might explain these observations.

There are well-known associations between vibration frequency, activated mechanoreceptors, and their receptive field sizes in different S1 subregions (Avanzino et al., 2014, Iwamura et al., 1993, Puckett et al., 2020, Weerakkody et al., 2007), which could partly explain the different results between low-(20Hz) and high-frequency (80Hz) stimulations. While the human back generally shows low tactile innervation density (Corniani and Saal, 2020), superficially located Meissner corpuscles (and Merkel disk receptors), which are preferentially activated by low-frequency stimulations, have smaller receptive fields than deeply located Pacinian corpuscles, which are activated by high-frequency stimulations (Cobo et al., 2021, Fleming and Luo, 2013, Iwamura et al., 1993, LaMotte and Mountcastle, 1975, Sherrick et al., 1990, Talbot et al., 1968). Hence, the *segmental model* might have well captured the organizational structure of brain activity patterns differentiating between large receptive fields of Pacinian corpuscles activated at 80Hz. However, this model may not have been able to accurately reflect the more localized similarity structures resulting from activation of Meissner corpuscles through 20Hz stimulation.

Furthermore, 80Hz stimulation might also have affected the activity of paraspinal muscle spindle afferents, the main proprioceptors of the human body (Goodwin et al., 1972, Proske and Gandevia, 2012, Roll et al., 1989). Therefore, one may assume that the cortical response to high-frequency stimulation reflects a mixture of different contributing mechanoreceptor types (deeper tactile and proprioceptive afferents with larger receptive fields), which might explain the broader informative brain activity patterns during 80Hz stimulation. This is further supported by the involvement of M1 regions (BA4a and BA4p), indicating that the central processing of paraspinal segmental afferent input involves a complex integration and sharing of back-specific topographic patterns across somatosensory and motor cortices.

### 4.2. The Importance of Cortical Topographic Mapping of the Back

Low back pain and movement behaviour are closely related and characterized by a tight interplay between the sensorimotor and pain processing systems at the spinal and supraspinal levels (Hodges and Tucker, 2011, Vittersø et al., 2022). Interest in the potential for a new therapeutic focus on sensorimotor integration is increasing (Bagg et al., 2022). One of the pain-relieving effects of such interventions is thought to be related to supraspinal processes, such as normalization of cortical sensory (proprioceptive) representations of the (lower) back through sensory discrimination training and graded exposure exercise (Van Dieën et al., 2019, Bagg et al., 2022). However. evidence that cortical representations of sensory afferents of the back are altered (“shifted” or “smudged”) in CLBP is sparse, and the pathological meaning of such potential alterations is unclear. In fact, only one study has identified a shifted representation towards the midline of somatosensory (tactile) input from the back in a small group of CLBP patients in S1 (Flor et al., 1997). The findings of that study, which used a magnetoencephalography technique to provide high temporal, but comparatively low spatial resolution recordings of brain activity, relative to fMRI, have never been replicated. Thus, the sensory-topographic maps of the human back that were revealed in the present study provide a sound basis for testing the “cortical map reorganization theory” in CLBP (Flor et al., 1997, Wand et al., 2011), which has recently been debated in other chronic pain conditions, such as complex regional pain syndrome (Mancini et al., 2019).

Furthermore, the observed involvement of the SPL in processing segmental paraspinal sensory input, a region lacking direct afferent projections (Passarelli et al., 2021), point to a diversified cortical organization of the back’s neural representation. SPL activations have been interpreted as encoding abstract movement representations (See et al., 2021). The functional connectivity of different SPL subregions, specifically of BA5 and BA7, to multiple functional networks, including salience, dorsal attention, and sensorimotor networks, has underscored the importance of the SPL in information processing along the attentional pathways (Alahmadi, 2021). Results from lesion studies in non-human primates and data from neurological patients have confirmed that the SPL is involved in coordinating spatial attention and motor accuracy (Passarelli et al., 2021). Hence, in the context of potential cortical reorganization of the back representation over time in pathologies such as CLBP, alterations in SPL activations might be indicative of wider implications when cortical topography is altered. Aside from a potential “de-differentiation” (Liu et al., 2021) of segmental paraspinal sensory input in S1, alterations in SPL organization might reflect the neural mechanism of the observed body schema disruptions of the trunk in CLBP patients (Bray and Moseley, 2011, Parkinson et al., 2010). However, the SPL has also been proposed to contain multiple subregions capable of flexible adaptation w.r.t. to coordination and motor control (Cunningham et al., 2013). Thus, it should be tested whether in CLPB patients compared to healthy controls such a flexible reorganization has taken place, which would be in line with the suggested “ reorganization theory” (Flor et al., 1997, Wand et al., 2011).

### 4.3. Limitations

Regarding potential muscle spindle activation, it is currently not possible to determine which specific muscle spindles are activated by the pneuVID. The primary assumption is that the superficial (longissimus and spinalis) muscles along the thoracolumbar axis are the most affected in terms of activation. Additional (rotatores and multifidi) muscles might be targeted, which show a particularly high density of muscle spindle fibers and are involved in key aspects of trunk and back proprioception (Boucher et al., 2015). Furthermore, it was not possible to completely isolate the contributions of superficial mechanoreceptors from those of the more deeply located mechanoreceptors. Precise anaesthetization of the skin could be used in future studies. This would allow to gauge the relative importance and topographical organization of proprioceptive afferents along the thoracolumbar axis as well as their cortical targets in more detail.

## Data and code availability

https://github.com/ISR-lab/RSA_HC

## Acknowledgements

The authors wish to thank Magdalena Suter for helping with data collection and recruitment of participants.

## References

Alexandre Abraham, Fabian Pedregosa, Michael Eickenberg, Philippe Gervais, Andreas Mueller, Jean Kossaifi, Alexandre Gramfort, Bertrand Thirion, and Gaël Varoquaux. Machine learning for neuroimaging with scikit-learn. Frontiers in neuroinformatics, page 14, 2014.

Adnan AS Alahmadi. Investigating the sub-regions of the superior parietal cortex using functional magnetic resonance imaging connectivity. Insights into Imaging, 12(1):1–12, 2021.

Laura Avanzino, Elisa Pelosin, Giovanni Abbruzzese, Michela Bassolino, Thierry Pozzo, and Marco Bove. Shaping motor cortex plasticity through proprioception. Cerebral Cortex, 24(10):2807–2814, 2014.

Matthew K Bagg, Benedict M Wand, Aidan G Cashin, Hopin Lee, Markus Hübscher, Tasha R Stanton, Neil E O’Connell, Edel T O’Hagan, Rodrigo RN Rizzo, Michael A Wewege, et al. Effect of graded sensorimotor retraining on pain intensity in patients with chronic low back pain: a randomized clinical trial. JAMA, 328(5):430–439, 2022.

Julien Besle, Rosa-Maria Sánchez-Panchuelo, Richard Bowtell, Susan Francis, and Denis Schluppeck. Event-related fMRI at 7T reveals overlapping cortical representations for adjacent fingertips in S1 of individual subjects. Human brain mapping, 35(5):2027–2043, 2014.

M Biggio, A Bisio, F Garbarini, and Marco Bove. Bimanual coupling effect during a proprioceptive stimulation. Scientific Reports, 11(1):15015, 2021.

Jean-Alexandre Boucher, Jacques Abboud, François Nougarou, Martin C Normand, and Martin Descarreaux. The effects of vibration and muscle fatigue on trunk sensorimotor control in low back pain patients. PloS one, 10(8):e0135838, 2015.

Helen Bray and G Lorimer Moseley. Disrupted working body schema of the trunk in people with back pain. British Journal of Sports Medicine, 45(3):168–173, 2011.

Matthew Brett, Chris Markiewicz, Michael Hanke, Marc-Alexandre Côté, Ben Cipollini, Paul McCarthy, Chris Cheng, Yaroslav Halchenko, Satra Ghosh, Eric Larson, Demian Wassermann, Stephan Gerhard, and Ross Markello. NiBabel, 2022.

Simon Brumagne, Martin Diers, Lieven Danneels, G Lorimer Moseley, and Paul W Hodges. Neuroplasticity of sensorimotor control in low back pain. journal of orthopaedic & sports physical therapy, 49(6):402–414, 2019.

Luke Chang, Sam, Eshin Jolly, Jin Hyun Cheong, Anton Burnashev, Andy Chen, Marissa Clar, Seth Frey, and Paxton Fitzpatrick. NLTools, 2021.

Yoon Gi Chung, Junsuk Kim, Sang Woo Han, Hyung-Sik Kim, Mi Hyun Choi, Soon-Cheol Chung, Jang-Yeon Park, and Sung-Phil Kim. Frequency-dependent patterns of somatosensory cortical responses to vibrotactile stimulation in humans: A fMRI study. Brain research, 1504:47–57, 2013.

Ramón Cobo, Jorge García-Piqueras, Juan Cobo, and José A Vega. The human cutaneous sensory corpuscles: an update. Journal of Clinical Medicine, 10(2): 227, 2021.

David M Cole, Philipp Stämpfli, Robert Gandia, Louis Schibli, Sandro Gantner, Philipp Schuetz, and Michael L Meier. In the back of your mind: Cortical mapping of paraspinal afferent inputs. Human Brain Mapping, 43(16):4943–4953, 2022.

Giulia Corniani and Hannes P Saal. Tactile innervation densities across the whole body. Journal of Neurophysiology, 124(4):1229–1240, 2020.

David A Cunningham, Andre Machado, Guang H Yue, Jim R Carey, and Ela B Plow. Functional somatotopy revealed across multiple cortical regions using a model of complex motor task. Brain research, 1531:25–36, 2013.

Halle R Dimsdale-Zucker and Charan Ranganath. Representational similarity analyses: a practical guide for functional MRI applications. In *Handbook of behavioral neuroscience*, volume 28, pages 509–525. Elsevier, 2018.

Halle R Dimsdale-Zucker, Maureen Ritchey, Arne D Ekstrom, Andrew P Yonelinas, and Charan Ranganath. CA1 and CA3 differentially support spontaneous retrieval of episodic contexts within human hippocampal subfields. Nature communications, 9(1):1–8, 2018.

Mary Beth Downs and Cindy Laporte. Conflicting dermatome maps: educational and clinical implications. journal of orthopaedic & sports physical therapy, 41(6): 427–434, 2011.

Naveed Ejaz, Masashi Hamada, and Jörn Diedrichsen. Hand use predicts the structure of representations in sensorimotor cortex. Nature neuroscience, 18(7):1034– 1040, 2015.

Joset A Etzel, Jeffrey M Zacks, and Todd S Braver. Searchlight analysis: promise, pitfalls, and potential. Neuroimage, 78:261–269, 2013.

Michael S Fleming and Wenqin Luo. The anatomy, function, and development of mammalian A*β* low-threshold mechanoreceptors. Frontiers in biology, 8:408–420, 2013.

Herta Flor, Christoph Braun, Thomas Elbert, and Niels Birbaumer. Extensive reorganization of primary somatosensory cortex in chronic back pain patients. Neuroscience letters, 224(1):5–8, 1997.

Guy M Goodwin, D Ian McCloskey, and Peter BC Matthews. Proprioceptive illusions induced by muscle vibration: contribution by muscle spindles to perception? Science, 175(4028):1382–1384, 1972.

Nina Goossens, Lotte Janssens, Madelon Pijnenburg, Karen Caeyenberghs, Char lotte Van Rompuy, Paul Meugens, Stefan Sunaert, and Simon Brumagne. Test– retest reliability and concurrent validity of an FMRI-compatible pneumatic vibrator to stimulate muscle proprioceptors. Multisensory Research, 29(4-5):465–492, 2016.

Gregory S Harrington and J Hunter Downs III. FMRI mapping of the somatosensory cortex with vibratory stimuli: Is there a dependency on stimulus frequency? Brain research, 897(1-2):188–192, 2001.

Paul W Hodges and Kylie Tucker. Moving differently in pain: a new theory to explain the adaptation to pain. Pain, 152(3):S90–S98, 2011.

Yoshiaki Iwamura, Michio Tanaka, Masahiro Sakamoto, and Okihide Hikosaka. Rostrocaudal gradients in the neuronal receptive field complexity in the finger region of the alert monkey’s postcentral gyrus. Experimental Brain Research, 92 (3):360–368, 1993.

Luke C Jenkins, Wei-Ju Chang, Valentina Buscemi, Matthew Liston, Peter Hum burg, Michael Nicholas, Thomas Graven-Nielsen, Paul W Hodges, James H McAuley, and Siobhan M Schabrun. Cortical function and sensorimotor plasticity are prognostic factors associated with future low back pain after an acute episode: the UPWaRD prospective cohort study. Pain, pages 10–1097, 2022.

Project Jupyter. Jupyter Notebook, 2021.

Eric R Kandel, James H Schwartz, Thomas M Jessell, Steaven A Siegelbaum, and AJ Hudspeth. Principles of neural science, fifth editon. In Principles of Neural Science. McGraw-Hill Education, 2013.

MG Kendall. Rank correlation methods 4th edition charles griffin. *High Wycombe*, Bucks, 1970.

Junsuk Kim, Yoon Gi Chung, Soon-Cheol Chung, Heinrich H Bülthoff, and Sung Phil Kim. Neural categorization of vibrotactile frequency in flutter and vibration stimulations: an fMRI study. IEEE transactions on haptics, 9(4):455–464, 2016.

James Kolasinski, Tamar R Makin, Saad Jbabdi, Stuart Clare, Charlotte J Stagg, and Heidi Johansen-Berg. Investigating the stability of fine-grain digit somatotopy in individual human participants. Journal of Neuroscience, 36(4):1113–1127, 2016.

Nikolaus Kriegeskorte, Rainer Goebel, and Peter Bandettini. Information-based functional brain mapping. Proceedings of the National Academy of Sciences, 103 (10):3863–3868, 2006.

Nikolaus Kriegeskorte, Marieke Mur, and Peter A Bandettini. Representational similarity analysis-connecting the branches of systems neuroscience. Frontiers in systems neuroscience, page 4, 2008.

Nikolaus Kriegeskorte, Jörn Diedrichsen, Marieke Mur, and Ian Charest. rsatoolbox, 2021.

Robert H LaMotte and Vernon B Mountcastle. Capacities of humans and mon keys to discriminate vibratory stimuli of different frequency and amplitude: a correlation between neural events and psychological measurements. Journal of Neurophysiology, 38(3):539–559, 1975.

Peng Liu, Anastasia Chrysidou, Juliane Doehler, Martin N Hebart, Thomas Wolbers, and Esther Kuehn. The organizational principles of de-differentiated topographic maps in somatosensory cortex. Elife, 10:e60090, 2021.

Chris Maher and Giovanni Ferreira. Time to reconsider what Global Burden of Disease studies really tell us about low back pain. Annals of the Rheumatic Diseases, 81(3):306–308, 2022.

Chris Maher, Martin Underwood, and Rachelle Buchbinder. Non-specific low back pain. The Lancet, 389(10070):736–747, 2017.

Tamar R Makin and Herta Flor. Brain (re) organisation following amputation: Implications for phantom limb pain. Neuroimage, 218:116943, 2020.

Flavia Mancini, Audrey P Wang, Mark M Schira, Zoey J Isherwood, James H McAuley, Giandomenico D Iannetti, Martin I Sereno, G Lorimer Moseley, and Caroline D Rae. Fine-grained mapping of cortical somatotopies in chronic complex regional pain syndrome. Journal of Neuroscience, 39(46):9185–9196, 2019.

Roberto Martuzzi, Wietske van der Zwaag, Juliane Farthouat, Rolf Gruetter, and Olaf Blanke. Human finger somatotopy in areas 3b, 1, and 2: a 7T fMRI study using a natural stimulus. Human brain mapping, 35(1):213–226, 2014.

Wes McKinney and the Pandas Development Team. Pandas (powerful Python data analysis toolkit, 2022.

Hamed Nili, Cai Wingfield, Alexander Walther, Li Su, William Marslen-Wilson, and Nikolaus Kriegeskorte. A toolbox for representational similarity analysis. PLoS computational biology, 10(4):e1003553, 2014.

Amy Parkinson, Laura Condon, and Stephen R Jackson. Parietal cortex coding of limb posture: in search of the body-schema. Neuropsychologia, 48(11):3228– 3234, 2010.

Lauretta Passarelli, Michela Gamberini, and Patrizia Fattori. The superior parietal lobule of primates: a sensory-motor hub for interaction with the environment. Journal of Integrative Neuroscience, 20(1):157–171, 2021.

Wilder Graves Penfield. Ferrier lecture-some observations on the cerebral cortex of man. Proceedings of the Royal Society of London. Series B-Biological Sciences, 134(876):329–347, 1947.

Haroon Popal, Yin Wang, and Ingrid R Olson. A guide to representational similarity analysis for social neuroscience. Social Cognitive and Affective Neuroscience, 14 (11):1243–1253, 2019.

Vencislav Popov, Markus Ostarek, and Caitlin Tenison. Practices and pitfalls in inferring neural representations. NeuroImage, 174:340–351, 2018.

Uwe Proske and Simon C Gandevia. The proprioceptive senses: their roles in sig naling body shape, body position and movement, and muscle force. Physiological reviews, 2012.

Alexander M Puckett, Saskia Bollmann, Keerat Junday, Markus Barth, and Ross Cunnington. Bayesian population receptive field modeling in human somatosen sory cortex. NeuroImage, 208:116465, 2020.

JP Roll, JP Vedel, and E Ribot. Alteration of proprioceptive messages induced by tendon vibration in man: a microneurographic study. Experimental brain research, 76(1):213–222, 1989.

W Schellekens, M Thio, S Badde, J Winawer, N Ramsey, and N Petridou. A touch of hierarchy: population receptive fields reveal fingertip integration in Brodmann areas in human primary somatosensory cortex. Brain Structure and Function, 226 (7):2099–2112, 2021.

Louis Schibli, Robert Gandia, Roger Buck, Philipp Staempfli, Michael Meier, and Philipp Schuetz. MR-safe multisegmental vibration device for cortical mapping of paraspinal afferent input. TechRxiv, 2021.

Geoffrey D Schott. Penfield’s homunculus: a note on cerebral cartography. Journal of neurology, neurosurgery, and psychiatry, 56(4):329, 1993.

Kyle B See, David J Arpin, David E Vaillancourt, Ruogu Fang, and Stephen A Coombes. Unraveling somatotopic organization in the human brain using machine learning and adaptive supervoxel-based parcellations. NeuroImage, 245:118710, 2021.

Carl E Sherrick, Roger W Cholewiak, and Amy A Collins. The localization of low and high-frequency vibrotactile stimuli. The Journal of the Acoustical Society of America, 88(1):169–179, 1990.

Stephen M Smith and Thomas E Nichols. Threshold-free cluster enhancement: ad dressing problems of smoothing, threshold dependence and localisation in cluster inference. Neuroimage, 44(1):83–98, 2009.

William H Talbot, Ian Darian-Smith, Hans H Kornhuber, and Vernon B Mountcastle. The sense of flutter-vibration: comparison of the human capacity with response patterns of mechanoreceptive afferents from the monkey hand. Journal of neurophysiology, 31(2):301–334, 1968.

Jaap H van Dieën, Herta Flor, and Paul W Hodges. Low-back pain patients learn to adapt motor behavior with adverse secondary consequences. Exercise and sport sciences reviews, 45(4):223–229, 2017.

Jaap H Van Dieën, N Peter Reeves, Greg Kawchuk, Linda R Van Dillen, and Paul W Hodges. Motor control changes in low back pain: divergence in presentations and mechanisms. Journal of Orthopaedic & Sports Physical Therapy, 49(6):370–379, 2019.

Axel D Vittersø, Monika Halicka, Gavin Buckingham, Michael J Proulx, and Janet H Bultitude. The sensorimotor theory of pathological pain revisited. Neuroscience & Biobehavioral Reviews, page 104735, 2022.

David A Walker. JMASM9: converting Kendall’s tau for correlational or meta analytic analyses. Journal of Modern Applied Statistical Methods, 2(2):26, 2003.

Alexander Walther, Hamed Nili, Naveed Ejaz, Arjen Alink, Nikolaus Kriegeskorte, and Jörn Diedrichsen. Reliability of dissimilarity measures for multi-voxel pattern analysis. Neuroimage, 137:188–200, 2016.

Benedict Martin Wand, Luke Parkitny, Neil Edward O’Connell, Hannu Luomajoki, James Henry McAuley, Michael Thacker, and G Lorimer Moseley. Cortical changes in chronic low back pain: current state of the art and implications for clinical practice. Manual therapy, 16(1):15–20, 2011.

Michael L Waskom. Seaborn: statistical data visualization. Journal of Open Source Software, 6(60):3021, 2021.

NS Weerakkody, DA Mahns, JL Taylor, and SC Gandevia. Impairment of human proprioception by high-frequency cutaneous vibration. The Journal of physiology, 581(3):971–980, 2007.

WR Willoughby, Kristina Thoenes, and Mark Bolding. Somatotopic arrangement of the human primary somatosensory cortex derived from functional magnetic resonance imaging. Frontiers in Neuroscience, 14:598482, 2021.

Kei Yamada, Yoshinari Nagakane, Kenji Yoshikawa, Osamu Kizu, Hirotoshi Ito, Takao Kubota, Kentaro Akazawa, Hiroyuki Oouchi, Shigenori Matsushima, Masanori Nakagawa, et al. Somatotopic organization of thalamocortical projection fibers as assessed with MR tractography. Radiology, 242(3):840–845, 2007.

